# HDAC6 promotes PMA-induced megakaryocyte differentiation of K562 cells by regulating ROS levels *via* NOX4 and repressing Glycophorin A

**DOI:** 10.1101/578096

**Authors:** Githavani Kummari, Ravi K Gutti, Arunasree M. Kalle

## Abstract

The human erythroleukemia (K562) cells are considered as bipotent megakaryocyte-erythroid progenitor cells and the differentiation of these cells to megakaryocytes (MK) in the presence of phorbol 12-myristate 13-acetate (PMA) mimics *in vivo* differentiation of MEP (megakaryocyte-erythroid progenitor) cells in the bone marrow. Histone deacetylases (HDACs) are involved in gene suppression and their roles during the MK differentiation remains largely undefined. In the present study, we have studied the expression levels of class I and class II HDACs during phorbol 12-myristate 13-acetate (PMA)-induced differentiation of K562 cells to MK. Class IIb HDACs (HDAC6 & HDAC10) were significantly up regulated time dependently upto 4 days of PMA-induced MK differentiation along with decreased acetylation levels of H3K9 and H3K56. Pharmacological inhibition and knockdown studies of HDAC6 using tubastatin A (TubA) and shRNA-HDAC6 respectively, during MK differentiation resulted in down regulation of MK lineage marker CD61 and up regulation of erythroid lineage gene glycophorin A (GYPA). HDAC6 over expression in K562 cells showed significant up regulation of CD61, MK transcription factors (FOG1 and GATA2) and down regulation of GYPA. ChIP-PCR studies showed enrichment of HDAC6 protein on GYPA promoter during differentiation indicating GYPA gene repression by HDAC6. Further studies on elucidating the role of HDAC6 in MK differentiation clearly indicated that HDAC6 is required for the production of sustainable levels of reactive oxygen species (ROS), an important regulator of MK differentiation, *via* NOX4.- ROS-HDAC6 circuit. In this study, we provide the first evidence that during PMA-induced megakaryocyte differentiation of K562 cells, HDAC6 represses erythroid lineage marker gene, GYPA, and promotes the sustainable levels of ROS *via* NOX4 required for MK differentiation.

**Key points:**

- HDAC6 upregulated during MK differentiation is involved in sustainable production of ROS *via* the circuit - HDAC6-NOX4-ROS-HDAC6.
- HDAC6 inhibits erythroid lineage gene, GYPA, by forming a repressor complex over the promoter region.

## Introduction

Survival, self-renewal and differentiation of hematopoietic stem cells in to blood cells is tightly regulated by expression of transcription factors, cytokines and miRNAs^1-5^. Megakaryocytes and erythrocytes are derived from a common megakaryocyte-erythroid progenitor (MEP) cells ^6^. Megakaryocytes (MK) produce enucleated platelets that are involved in blood clotting, hemostasis and in immune response ^7^. Thrombocytopenia, reduced platelets count, is a clinical problem in many disease conditions such as cancer therapy, trauma, sepsis and viral infections such as dengue, etc ^8^ and requires platelet transfusion. The two reasons for thrombocytopenia are reduced production from MK cells or increased destruction of platelets^9^. Better understanding of molecular mechanisms regulating MK differentiation and platelet production are required to identify the safe sources of platelets for therapeutic purposes and to cure the platelet diseases.

K562 cells are chronic myeloid leukemia cells, well studied model system to find out the molecular mechanisms regulating the erythroid and megakaryocyte lineages in the presence of different chemical inducers ^10^. PMA (Phorbal 12-myristate 13-acetate) induces the differentiation of K562 cells towards megakaryocyte lineage whereas hemin, hydroxyurea and Ara-C (Arabinosyl cytosine) induce erythroid differentiation of these cells ^11,12^. The differentiation of K562 cells to megakaryocytes is monitored by cell growth arrest, changes in morphology, endomitosis and adhesive properties due to expression of integrins CD61 and CD41 (MK markers)^13^ and reduced expression of erythroid genes Glycophorin A (GYPA). PMA induces the activation of PKC leading to further ERK1/2, c-JUN, c-FOS and NF-κB transcription factor activation ^14-17^. PMA also induces ROS production in K562 cells and ROS is an important factor driving MK cell differentiation ^18^.

The two epigenetic marks that play an important role in gene regulation are methylation and acetylation. Acetylation and deacetylation of histone proteins is controlled by histone acetyl transferases (HATs) and histone deacetylases (HDACs) respectively. Deacetylation of histone proteins by HDACs leads to the formation of compact chromatin which prevents the binding of transcriptional machinery and thus gene suppression ^19^. HDACs play important role in various physiological and pathological processes by regulating the activity of non-histone proteins by deacetylation ^20^. Total of 18 HDACs have been identified in humans and divided into four classes based on their homology to yeast orthologs. Class I (HDAC1,2,3,4,8), Class II (HDAC4,5,7,9 & 6,10) and class IV (HDAC11) are classical zinc dependent enzymes whereas class III HDACs are named as sirtuins from SIRT1-7 which requires NAD+ as cofactor for their activity ^21-23^. Recent studies indicate that the epigenetic gene regulation governs the stem cell differentiation and the role of HDAC inhibitors (HDACi) in reprogramming of somatic cells to pluripotency is well established ^24^.

The commitment of the bipotent progenitor cells to a particular lineage requires suppression of the opposite lineage genes. Since HDACs are involved in gene repression *via* compact chromatin formation, we hypothesize that one or few of the HDACs might be involved in lineage commitment of progenitor cells. Although the involvement of HDACs in hematopoiesis is known, the role of HDACs and the underlying molecular mechanisms of lineage commitment remain elusive. Recently, Messaoudi et al have shown that, HDAC6 plays an important role in proplatelet release from MKs by deacetylating cortactin, an important cytoskeletal protein^25^. We therefore, in the present study, aimed at identifying the role of HDACs in MK lineage commitment of K562 cells.

## Methods

### Cell culture

K562 cells were obtained from the National Centre for Cell Science (NCCS) cell repository, India. The cells were cultured in RPMI 1640 medium supplemented with 10% FBS (Fetal Bovine Serum) and 1% penicillin and streptomycin and grown at 37 °C, 5% CO_2_. The exponentially growing K562 cells were seeded at a density of 2.5 ×10^5^ cells / ml and treated with 20 nM PMA for 24 h and 48 h. In combination treatments, the cells were treated with 5 µM tubastatin A or 100 µM quercetin, 1 h prior to PMA addition and the control cells received DMSO alone.

### RNA isolation and RT-PCR

Total RNA was isolated by TRI reagent (Ambion) according to the manufacture’s protocol and quantification and purity of RNA were determined by Nanodrop. The integrity of RNA was examined by agarose gel electrophoresis. 2 µg of RNA was reverse transcribed with Oligo(dT)15, 50 mM Tris-HCl pH 8.3, 75 mM KCl, 3 mM MgCl2, 10 mM dithiothreitol, 200 μM each deoxynucleotide triphosphate [dNTP]) using 200 U M-MLV Reverse Transcriptase (Invitrogen) in 20 μl of a reaction mixture. Following denaturation at 65°C for 5 min, the reaction mixture was incubated at 42 °C for 1 h. Target genes were quantified by using SYBR green chemistry, gene specific primers with appropriate dilutions of cDNA. The expression levels were calculated using 2^-ΔΔCT^ method and normalized with control cells. The primers (Table1) were designed using UCSC Genome browser software. GAPDH was used as reference gene to calculate the changes in target gene expression levels.

**Table 1:**
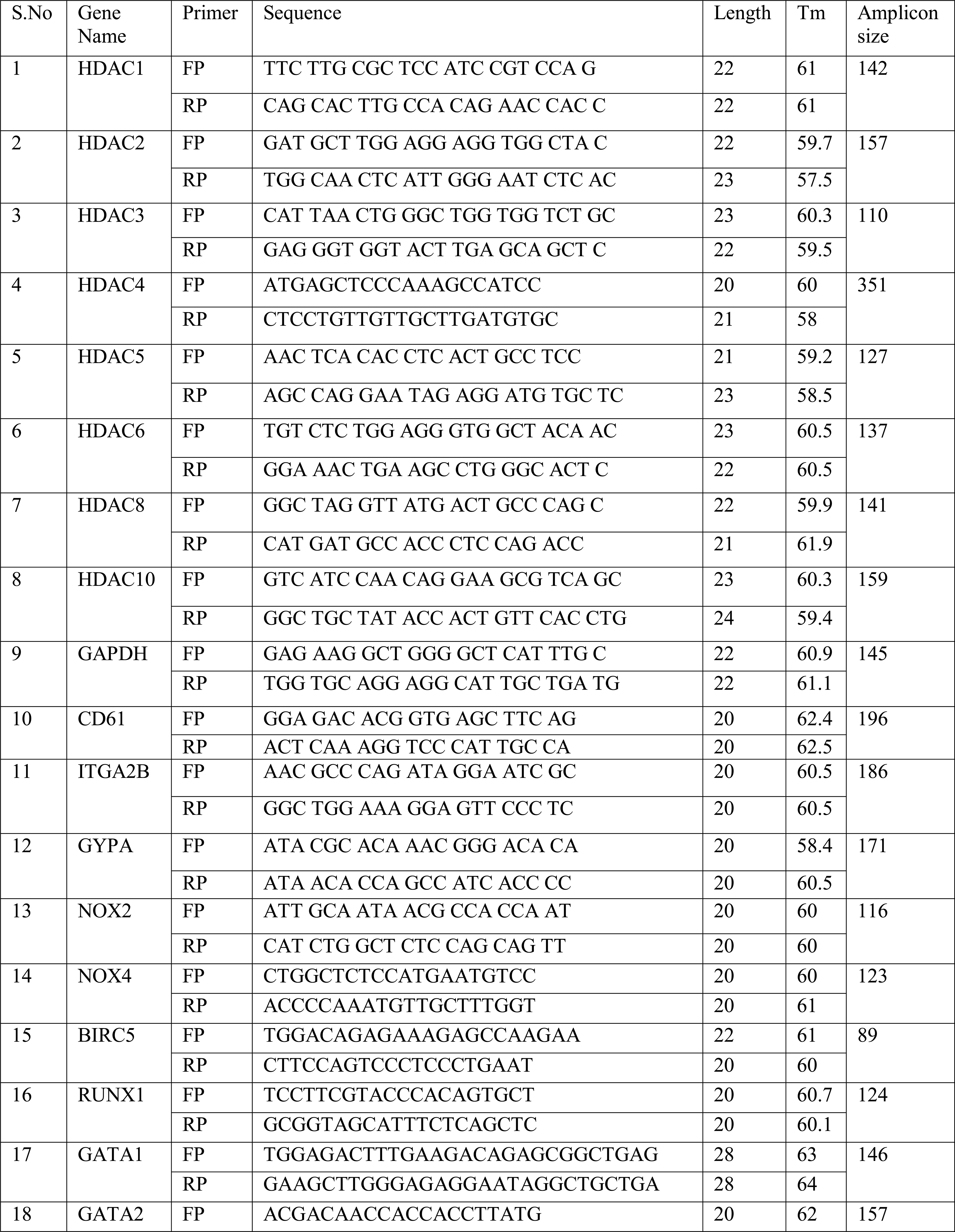
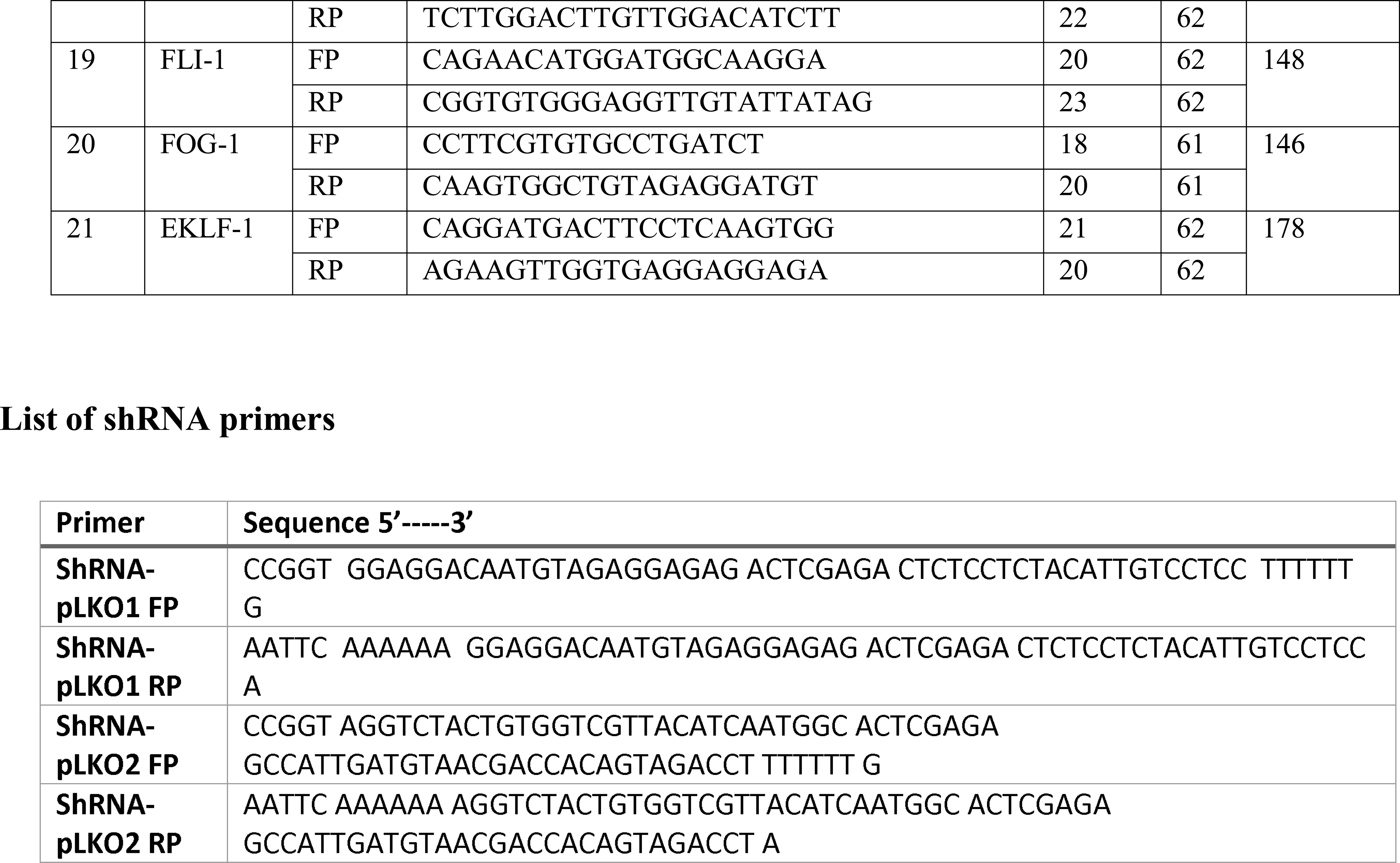
List of Primers

### Western blotting

Following different treatments for the times indicated in figures, the cells were lysed in RIPA buffer [20 mM Tris/HCl (pH 7.5), 150 mM NaCl, 1 mM sodium EDTA, 1 mM EGTA, 1%Triton, 2.5 mM sodium pyrophosphate, 1 mM 2-glycerophosphate, 1 mM sodium orthovanadate, 1 μg/ml leupeptin and 1 mM PMSF) on ice for 30 min. The cell lysates were then centrifuged at 13000 g for 30 min at 4 °C. The supernatant containing protein was quantified by BCA (bicinchoninic acid) assay (Santacruz). Equal amounts protein was separated on 10% SDS-PAGE and transferred on to nitrocellulose membrane. The membrane was blocked with 3% nonfat skimmed milk powder for 45 min. Specific proteins were probed with primary antibodies-HDAC6, HDAC10, H3, acetyl tubulin, tubulin, Ac-H3K9, Ac-H3K56 and GAPDH (Cell signaling). After washing the membrane with TBS-T for 3 times, each wash for 10 min; secondary antibody conjugated with horseradish peroxidase was added and incubated for an hour. After two washes with TBS-T for 10 min each, detection was performed by enhanced chemiluminiscence reagent (ECL, Amersham) and captured using BioRad imaging system.

### Isolation of Histone proteins

Histone proteins were isolated from treated and control cells as described in Nature protocols^26^. In brief, the cells were washed with ice cold PBS and resuspended in 5-10 volumes of hypotonic solution (10 mM Tris pH 8,10 mM KCl, 1.5 mM MgCl_2_ and 1 mM DTT) and incubated on ice for 1 h. Intact nuclei were pelleted at 10000 g, 4 °C for 10 min and then resuspended in acidic solution (0.4 N H_2_SO_4_, 10 mM HEPES, 1.5 mM MgCl_2_, 10 mM KCl, 0.5 mM DTT, 1.5 mM PMSF, and protease inhibitor cocktail) with mixing on rotospin for 30 min. The supernatant containing histone proteins collected at 16000 g at 4 °C for 10 min and precipitated by adding 132 µl of 100% TCA (Tri cholroacetic acid). After 30 min incubation on ice, the histone proteins were pelleted and washed with acetone twice to remove the acid. Finally the protein was air dried and dissolved in water. Immunoblotting technique was used to analyze the different acetylation levels of lysine residues of histone protein (H3) as mentioned above.

### Transfection

K562 cells (1 ×10^6^) were transfected with plasmid pcDNA-HDAC6-FLAG using PEI reagent as per the manufacture’s protocol. Briefly, the plasmid DNA was mixed with PEI reagent in the ratio of 1:1 in serum free media and incubated for 15 min before adding to the cells. After 3 h incubation, complete medium was added and allowed to grow for different time points. Real-time PCR, Immunoblot and HDAC activity assay were used to confirm the overexpression. pcDNA-HDAC6-FLAG was a gift from Dr.Tso-Pang Yao (Addgene plasmid # 30482).

### Nuclear and cytoplasmic extractions

Briefly, the cells were lysed in cytosolic extraction buffer (30 mM Tris-HCl, [pH 7.5], 150 mM NaCl, 10 mM MgCl_2_, 1% Nonidet P-40, 1 mM PMSF and protease inhibitor cocktail), incubated on ice for 3 min and centrifuged at 10000 g for 3 min. The pelleted nuclei were washed with cytosolic extraction buffer and spun at 16000 g for 5 min. The nuclei were lysed in nuclear extraction buffer (10 mM HEPES [pH 7.9], 420 mM NaCl, 10 mM MgCl_2_, 1 mM EDTA, 25% glycerol, 1 mM PMSF and protease inhibitor cocktail) for 30 min on ice with intermittent vortexing. The nuclear extract was collected at 16000 g for 30 min at 4°C.

### Immuno Fluorescence

The control and PMA treated cells were fixed in 4% formaldehyde, permeabilized with 0.25% Triton X-100 and blocked with 3% bovine serum albumin in phosphate-buffered saline. The cells were incubated with primary HDAC6 antibody for 1 h, followed by washes with 1%BSA in PBS-T and then the cells were incubated with fluorophore-conjugated secondary antibody (Alexa 647) at room temperature and 4, 6-diamidino-2-phenylindole was used to for nuclear staining. Mounted samples were viewed under a Carl Zeiss, NLO 710 laser scanning confocal microscope and images were captured with ZEN 2010 acquisition software.

### Measurement of ROS by Flow cytometry

2′, 7′-dichlorodihydrofluorescein diacetate (H_2_H2DCF-DA) is an oxidation sensitive dye and was used to measure the ROS production in cells treated with PMA or / and tubastatin A. The cells were washed with PBS and then incubated with 5 μM H_2_H2DCF-DA in PBS for 30 min at 37 °C. The cells were collected and analyzed using BD LSR Fortessa flow cytometry system.

### Chromatin Immuno precipitation (ChIP)

Cells (2.5 × 10^5^) were fixed in 1% formaldehyde solution in 5 ml of serum free media for 10 min at room temperature followed by quenching with 0.125 M glycine for 5 min. Then the cells were washed twice with PBS and resuspended in cell lysis buffer for 10 min. Nuclear pellet was resuspended in nuclear lysis buffer and IPDB (immune precipitation dilution buffer) and subjected to sonication at an amplitude of 36%, 25-30 cycles, each cycle with 20 sec on and 40 sec off for shearing the DNA into fragments of 300-1000 bp. Cell debris is removed by centrifugation at 14000 rpm for 5 min at 4°C. The chromatin was diluted with NLB: IPDB in 1:4 ratio and precleared with rabbit IgG. Then 100 µl of chromatin sample was used as input and the remaining was incubated with HDAC6 antibody (Abcam ab47181), for overnight at 4°C. Dynabeads® Protein G (Invitrogen 10004D) were used to collect the immune precipitated protein-DNA complexes. The DNA from input and IP samples was purified by phenol chloroform method and quantified using Nanodrop. PCR was performed to amplify GYPA promoter with the following primers sets. Set1 FP-5’ CTT GAG CAC AAT TCC TGC AA 3’, RP 5’ CTG AGC AGC AGG ACA AGA A 3’, Set2 FP5’ ATT GAG CTT CCT CGC ATT TT 3’, RP 5’ CGC AGC TAT GAA ACC AGT GA 3’ set3 FP 5’ GGC TCC ACA ACA GCT ACC TC 3’ RP5’ TCT TGG GGC TAT GAA AGT GG 3’, set4 FP 5’ AAA TGC CTC CCC TGC CTA T 3’ RP5’ CCT GAG ATC ATG AGC TGG TTC 3’set5 FP 5’ CAA GGG AGC CCA GTA TTT ATG 3’ RP5’ GCC ATT TGG CAG AAA TAG GA 3’ GAPDH primers FP 5’ TAC TAG CGG TTT TAC GGG CG 3’, RP 5’ TCG AAC AGG AGG AGC AGA GAG CGA 3’.

### Knockdown of HDAC6 expression using shRNA

The shRNA oligos were designed, cloned in pLKO.1 puro vector and confirmed as per the protocol from addgene (Addgene Plasmid 10878. Protocol Version 1.0. December 2006.) shRNA sequences GGAGGACAATGTAGAGGAGAG, AGGTCTACTGTGGTCGTTACATCAATGGC, targeting different regions of HDAC6, were chosen from Block it design tool and Origene company and ordered from Integrated DNA Technologies. pLKO.1 puro control shRNA was a generous gift from Dr. Kishore Parsa from DRILS, Hyderabad. Transfection was done as mentioned above.

### HDAC activity assay

HDAC activity was determined using fluorescent substrate FLUOR DE LYS® (ALX-260-137-M005) purchased from Enzo Life Sciences as per the manufacturer’s protocol. The HDAC6 protein was immunoprecipitated from total protein lysates (0.25 mg) of cells treated with different compounds using protein A agarose beads (Sigma). The HDAC6 protein bound beads were used as enzyme source and was incubated with the HDAC assay buffer (50 mM Tris pH 8.0, 137 mM NaCl,2.7 mM KCl, 1 mM MgCl2 and 1 mg/ml BSA) and fluorescent substrate (60 μM final) to a final volume of 50 μl at 37 °C for 15 min. Reaction was stopped with the addition of 50 μl developer solution (Enzo Life Sciences) followed by incubation at 37 °C for 30 min. The Arbitrary Fluorescent Units (AFU) was recorded at Excitation wavelength of 360 nm and Emission wavelength of 460 nm.

### Statistical analyses

Results were expressed as the means of ±standard deviation (SD) of at least three independent experiments. Statistical analysis was carried out by one-way ANOVA, two-way ANOVA, followed by Tukey’s multiple comparison tests using Graphpad Prism 6.01. The statistical significance is represented as *p-value < 0.05; **p-value < 0.01 and ***p-value < 0.001.

## Results

### Differential expression of classIIb HDACs with decreased acetylation levels of H3K9 and H3K56 during MK differentiation of K562 cells

PMA-induced MK differentiation of K562 cells was confirmed by change in morphology of cells (Fig 1A), upregulation of MK markers such as CD41 and CD61 and downregulation of an erythroid marker glycophorin A (GYPA) (Fig 1B). Once the model system was validated, we next did HDAC expression profiling (mRNA and protein expression). The expression levels of class I (HDAC1, 2 and 8) and class IIa HDACs (HDAC4) did not change significantly in control and PMA-treated cells (Fig 1C & D). However, RNA and protein levels of class IIb HDACs, HDAC6 and 10, were significantly upregulated during MK differentiation (Fig 1C & D). Further, time-course mRNA expression analysis indicated upregulation of HDAC6 along with MK markers till 4 days of PMA treatment and gradual downregulation of GYPA significantly (Fig. 1E). Since histones are the primary substrates for HDACs, we have analyzed the global acetylation levels of lysine residues of histone H3. We have observed a significant decrease in acetyl levels of H3K9 and H3K56 during PMA-induced differentiation of K562 cells (Fig 1F), suggesting these might be the targets of upregulated class IIb HDACs. The acetylation levels of H3K18, H3K14 and H3K27 did not alter in control and PMA treated cells (Fig 1F).

**Fig 1:**
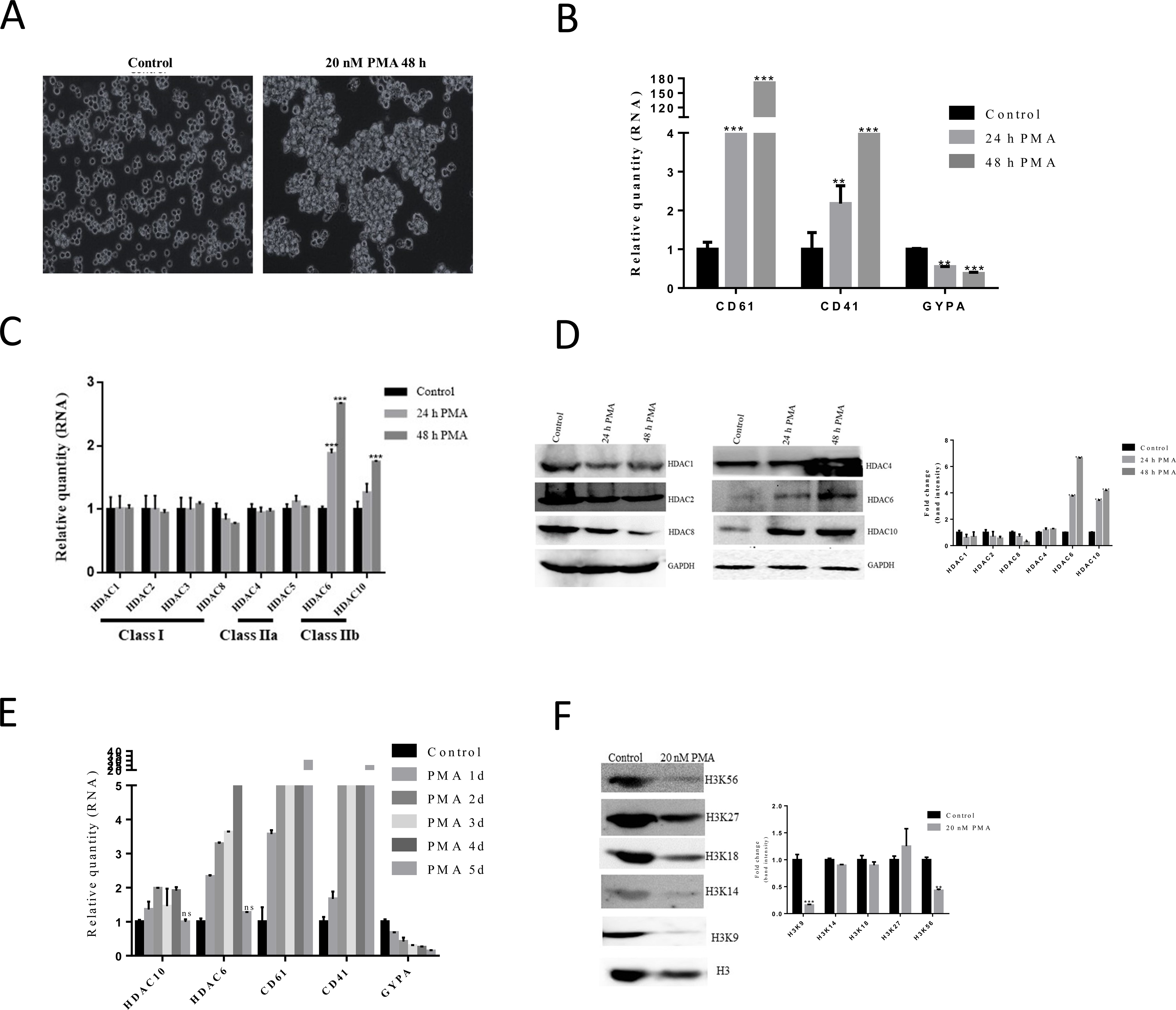
Differential expression of classIIb HDACs with decreased acetylation levels of H3K9 and H3K56 during MK differentiation of K562 cells. Phase contrast microscopy of K562 cells treated with PMA (A), RNA expression levels of MK lineage markers CD61 and CD41 and erythroid gene(GYPA) in PMA treated cells at different time points (B), RNA expression levels of class I and II HDACs (C), protein expression of HDAC1, 2, 8, 4, 6 and 10 during MK differentiation and densitometry by Image J software (D), RNA expression of HDAC6, 10 and MK markers and GYPA in PMA treated cells for 5 days (E), Western blotting of acetylation levels of different lysine residues of H3 protein in control and PMA treated cells and densitometry by Image J software (F).

### HDAC6 upregulation is specific to K562 cell differentiation to MK cells

PMA is known to activate several signaling pathways including PKC and PI3K. So we asked the question whether HDAC6 upregulation is due to PMA signaling or is MK lineage specific. To address this, we assessed the expression levels of HDAC6 in PMA-induced HL-60 cells that differentiate into monocytes. The results showed no change in HDAC6 expression levels in PMA-induced HL-60 cells indicating that HDAC6 upregulation is MK lineage specific (Fig. 2A). To better understand the role of HDAC6 in MK differentiation as a repressor of gene or as a protein activity regulator, we first investigated HDAC6 localization by confocal microscopy. As shown in Fig. 2B, HDAC6 is predominantly localized to the nucleus during MK differentiation when compared to pan-cellular distribution in control cells. Immunoblot analysis of the cytoplasmic and nuclear fractions also indicated similar results (Supplementary Fig. S1). To get an insight into the nuclear function of HDAC6, we analyzed the acetylation levels of the two histone H3 lysine residues, H3K9 and H3K56, that were actually downregulated during MK differentiation (Fig. 1F) in presence of HDAC6-specific inhibitor, Tubastatin A (Tub A). The specificity of Tub A towards HDAC6 was confirmed by analyzing the Ac-tubulin levels dose-dependently (Supplementary Fig. S2) and also its inhibitory effect on other HDACs by Immunoblot (Supplementary Fig. S3). The Ac- H3K9 and Ac-H3K56 immunoblot showed a significant increase in the acetylation levels of H3K9 and H3K56 (Fig. 2C) suggesting that HDAC6 might be playing an important role as a transcriptional regulator during PMA-induced differentiation of K562 cells to MK cells. We next investigated the mechanism by which HDAC6 is upregulated during MK differentiation of K562 cells. We analyzed the PKC pathway activated by PMA in K562 cells during MK differentiation. We determined the levels of phospho-ERK and Ac-NF-κB in control and PMA-treated K562 cells. In the immunoblots, we found increased p-ERK levels and increased Ac-NF-κB (K310) along with increased HDAC6 during PMA treatment (Fig 2D). Further, we found reduced HDAC6 expression when ERK1/2 activation was inhibited with apigenin (Fig 2E) during PMA induced differentiation of K562 cells suggesting that HDAC6 upregulation during MK differentiation is *via* PKC-p-ERK-NF-κB axis.

**Fig 2:**
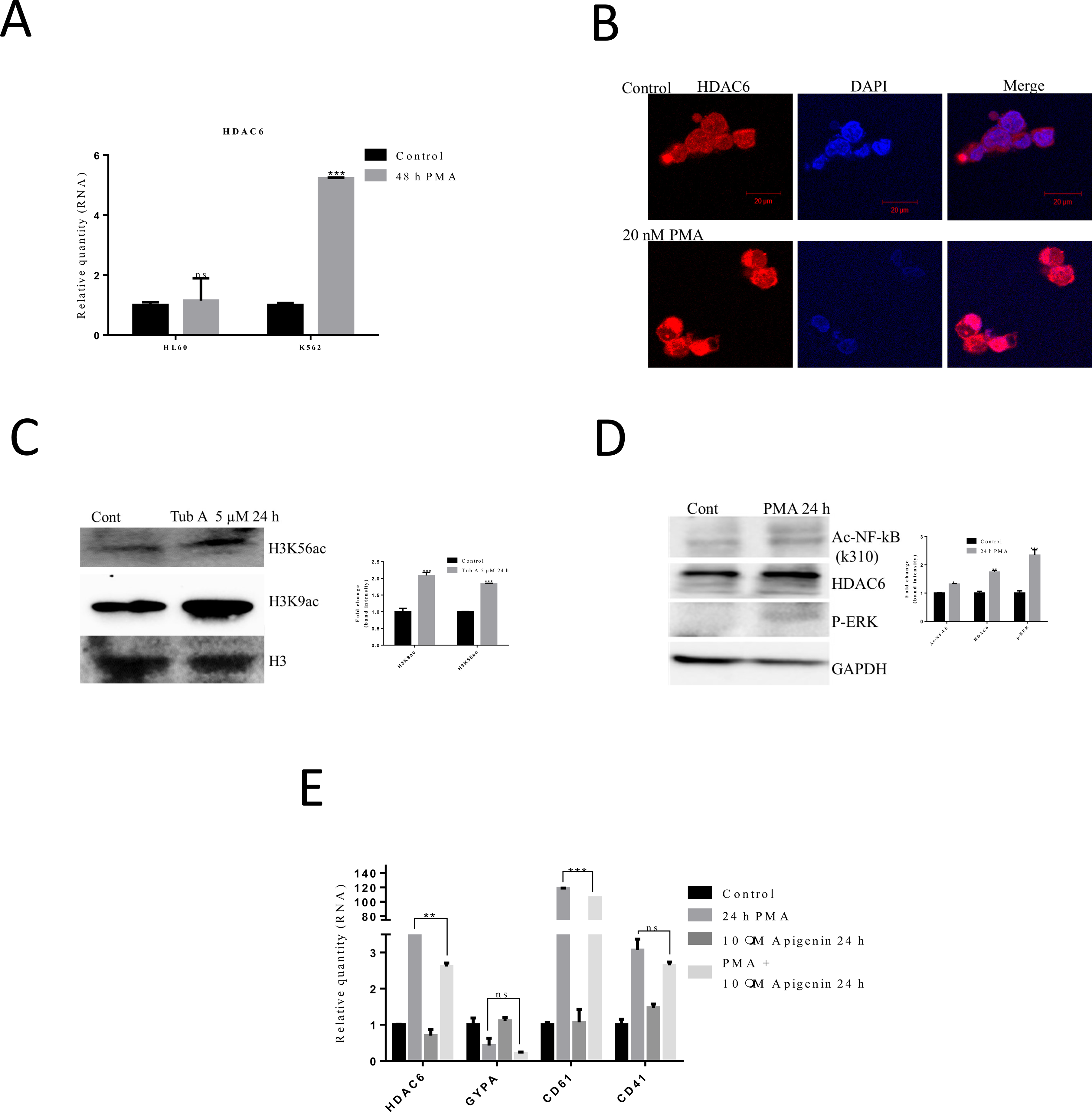
HDAC6 upregulation is specific to K562 cell differentiation to MK cells. HDAC6 expression in HL-60, K562 cells treated with 20 nM PMA for 48 h(A), localization of HDAC6 in control and PMA treated cells by confocal microscopy (B). acetylation levels of H3K9 and H3K56 in the presence of HDAC6 inhibitor and densitometry by Image J software (C), Western blotting of HDAC6, ac-NF-kB, p-ERK in control and PMA treated cells and densitometry by Image J software (D), RNA expression analysis of HDAC6, MK markers, GYPA in ERK1/2 inhibitor (Apigenin) treated cells duing MK differentiation (E).

### HDAC6 is required for the expression of MK markers and inhibition of erythroid marker

To further delineate the role of HDCA6 during MK differentiation, we have treated the cells with Tub A, and analyzed the expression levels of MK lineage genes. The qPCR results clearly indicated that MK markers were downregulated when HDAC6 is inhibited in PMA treated cells along with significant upregulation of erythroid marker (Fig 3A). We also measured the protein levels of CD61 by flow cytometry and the results are in agreement with qPCR data (Fig 3B (i) & (ii)). Next we performed HDAC6 knockdown during MK differentiation with shRNA (pLKO.1 HDAC6) and studied the MK linage commitment. The expression analysis of MK marker (CD61) and erythroid marker (GYPA) showed significant reduced levels of CD61 with significant induction of GYPA during MK differentiation (Fig 3C), mimicking effects obtained with Tub A. To further probe the effect of HDAC6 during MK differentiation, we have overexpressed HDAC6 in K562 cells and studied the MK lineage marker expression. Overexpression of HDAC6 in K562 cells was confirmed at the RNA level by qPCR (Fig 3D (i)), protein level by immunoblot (Fig 3D (ii)) and activity by HDAC assay (Fig 3D (iii)). Overexpression of HDAC6 resulted in significant upregulation of CD61, but not CD41 and downregulation of erythroid lineage gene, GYPA (Fig 3E). These results suggest that HDAC6 might be negatively regulating erythroid lineage gene, GYPA expression and positively regulating the MK lineage gene, CD61 expression. We then sought to determine how HDAC6 is positively regulating the expression of MK marker (CD61). To explore this, we have analysed the expression levels of MK transcription factors GATA2, FOG-1 and GATA1 in HDAC6 inhibited cells treated with PMA. We observed reduced expression of GATA2 and FOG-1 in Tub A treated cells alone but not in HDAC6 inhibited cells treated with PMA (Fig 3F). On the contrary, we observed upregulation of GATA2 and FOG-1 in HDAC6 overexpressed cells without PMA treatment (Fig 3G). These results suggest that HDAC6 might be regulating the expression of GATA2 and FOG-1 indirectly that in turn regulate CD61 expression.

**Fig 3:**
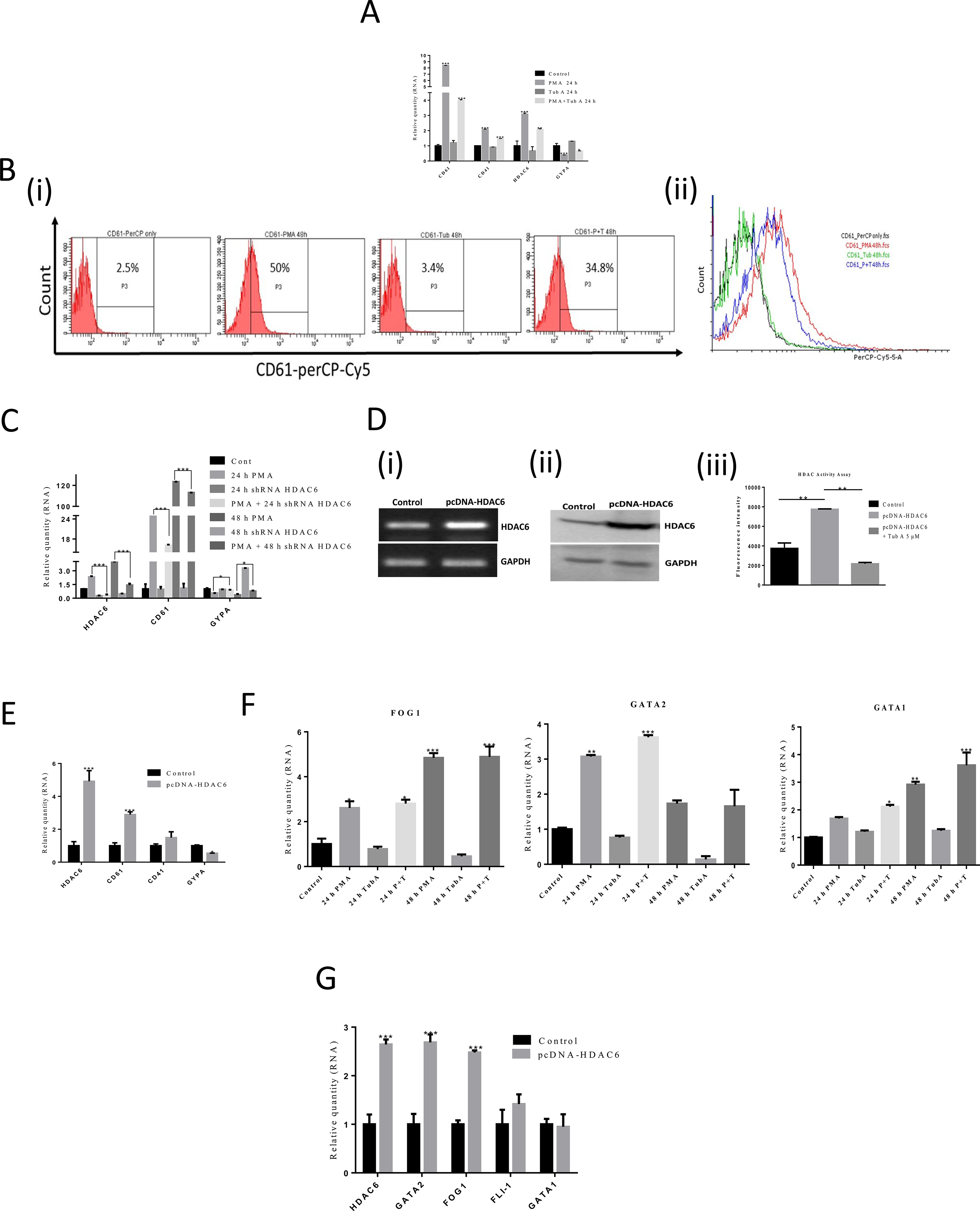
HDAC6 is required for the expression of MK markers and inhibition of erythroid marker. RNA expression levels of MK markers(CD61, CD41), HDAC6 and GYPA in cells treated with PMA and or tubastatin A for 24 h (A), CD61 protein expression by flow cytometry in cells treated with PMA and or tubastatin A for 48 h (B(i)) and overlay of histograms (B(ii)), RNA expression analysis of HDAC6, CD61 and GYPA in HDAC6 knockdown cells for 24 h and 48 h in the presence or absence of PMA (C), RNA expression analysis of CD61, CD41 and GYPA in HDAC6 transfected cells(D(i)), HDAC activity assay (D(ii)), transcription factor expression in HDAC6 inhibited (E), and overexpressed cells (F).

### HDAC6 represses GYPA promoter

When HDAC6 was inhibited with tubastatin A, we have observed upregulation of GYPA not only in PMA-treated K562 cells, but also in K562 cells treated with Tub A alone (Fig 4A). But we did not observe significant effect on other erythroid lineage genes like γglobin and EKLF (Fig 4B). This prompted us to look into the role of HDAC6 in GYPA transcriptional repression. We therefore designed 5 sets of primers from first intron to the 1400 bp upstream to the transcription start site (TSS) of GYPA promoter (Fig. 4C) and did ChIP-PCR with HDAC6 antibody. We observed a 2-fold enrichment of HDAC6 over the GYPA promoter at 4 out of 5 sites studied in control cells (Fig 4D). However, there was a significant enrichment at site 1 (−1363 bp to 1183 bp) in PMA-treated cells. The HDAC6 binding decreased significantly at site 1 when treated with Tub A (Fig 4D) in control cells. These results clearly indicated that HDAC6 plays an important role in repressing erythroid-specific lineage gene, GYPA.

**Fig 4:**
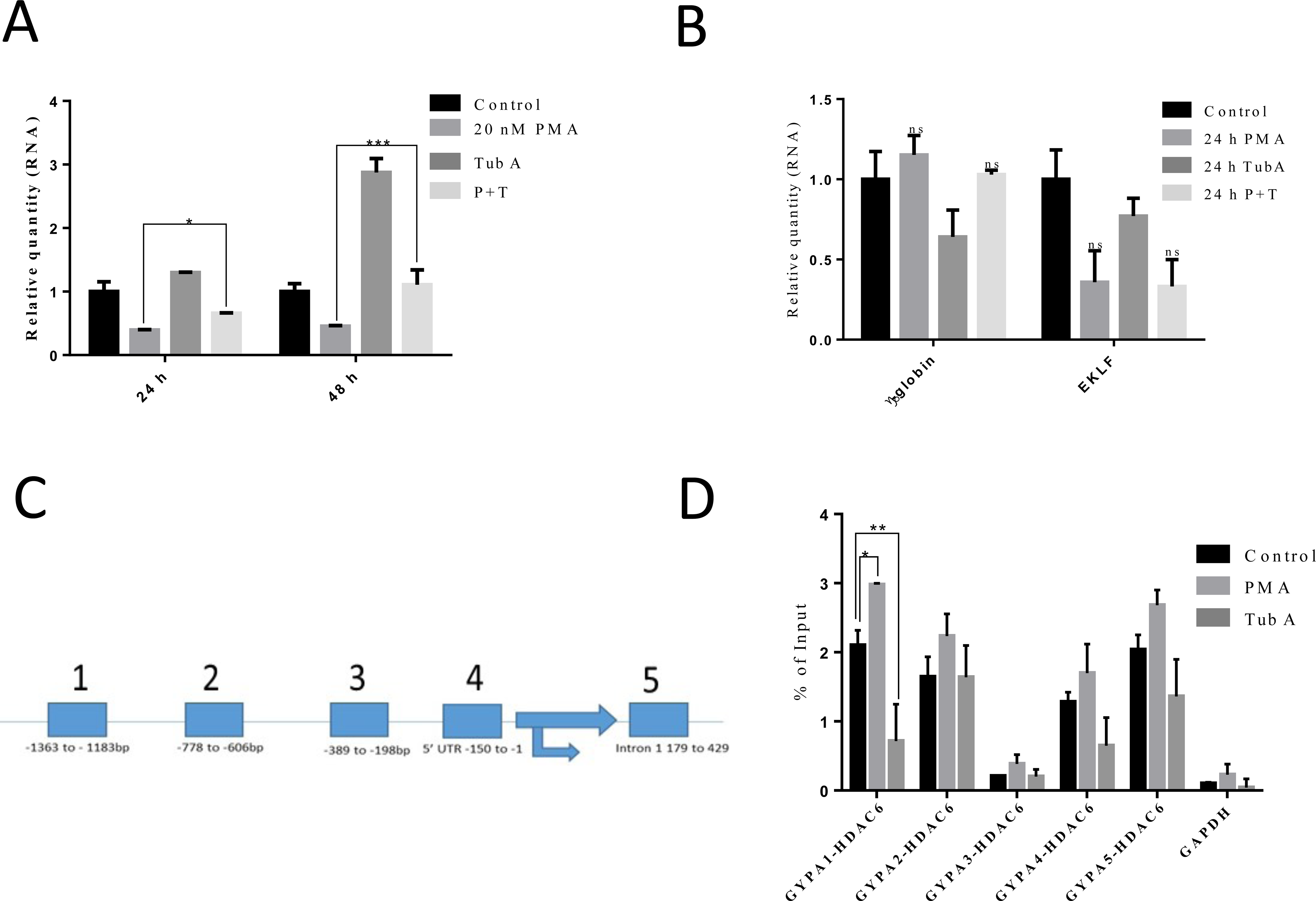
HDAC6 represses GYPA promoter. RNA expression levels of GYPA in PMA and / or TubA treated cells (A), RNA expression levels of globin and EKLF in PMA and / or TubA treated cells (B), 5 sets of primers designed from first intron to the 1400 bp upstream to the transcription start site (TSS) of GYPA promoter (C), ChIP-PCR with HDAC6 for different sites of GYPA promoter using specific primers in PMA or tubastatinA treated cells (D).

### Crosstalk between PMA signalling, ROS and HDAC6 during MK differentiation

The strong nuclear localization of HDAC6 in the present study correlated, partly, with the GYPA gene repressing function of HDAC6. However, the immunofluorescence and immunoblot data indicated cytoplasmic presence of HDAC6. So we also focused on other probable functions HDAC6 might be having during MK differentiation. Since the role of HDAC6 in ROS production and the role of ROS in MK biology is well known, we tried to find a link between ROS and HDAC6 in MK differentiation. We measured ROS levels using H2DCF-DA by flow cytometry and observed increased ROS levels in PMA treated cells that were reduced upon HDAC6 inhibition (Fig 5A). Since NADPH oxidases are involved in ROS production, we next examined the effect of Tub A on NOX2 and NOX4 expression and found that NOX4 was upregulated during MK differentiation and significantly downregulated in the presence of HDAC6 inhibitor with no change in NOX2 expression (Fig 5B) suggesting for the first time that PMA-induced ROS production in MK differentiation of K562 cells is *via* NOX4. In addition, we have observed downregulation of NOX4 in HDAC6 knockdown cells (Fig 5C). To further understand the relation between HDAC6 and NOX4, we analyzed the effect of an antioxidant, quercetin, on the expression of HDAC6, NOX2, NOX4 and MK markers during PMA-induced differentiation of K562 cells. During MK differentiation in the presence of antioxidant, significant reduction in the expression levels of HDAC6 and NOX4 along with MK markers was observed (Fig 5D). This result implicated a clear crosstalk between HDAC6, NOX4 and ROS during MK differentiation. Next, we confirmed the role of HDAC6 in ROS production via NOX4 in MK biology, by analysing the mRNA levels of survivin gene, a chromosome passenger protein involved in cell division. During MK differentiation, survivin is downregulated so as to promote polyploidy in MK cell. We observed that survivin is upregulated when HDAC6 is inhibited (Fig 5E) suggesting that ROS produced by NOX4 is important for polyploidy of MK cell and that HDAC6 is involved in ROS production. Taken together, these results confirm the interplay between ROS, NOX4 and HDAC6 during PMA induced differentiation of K562 cells.

**Fig 5:**
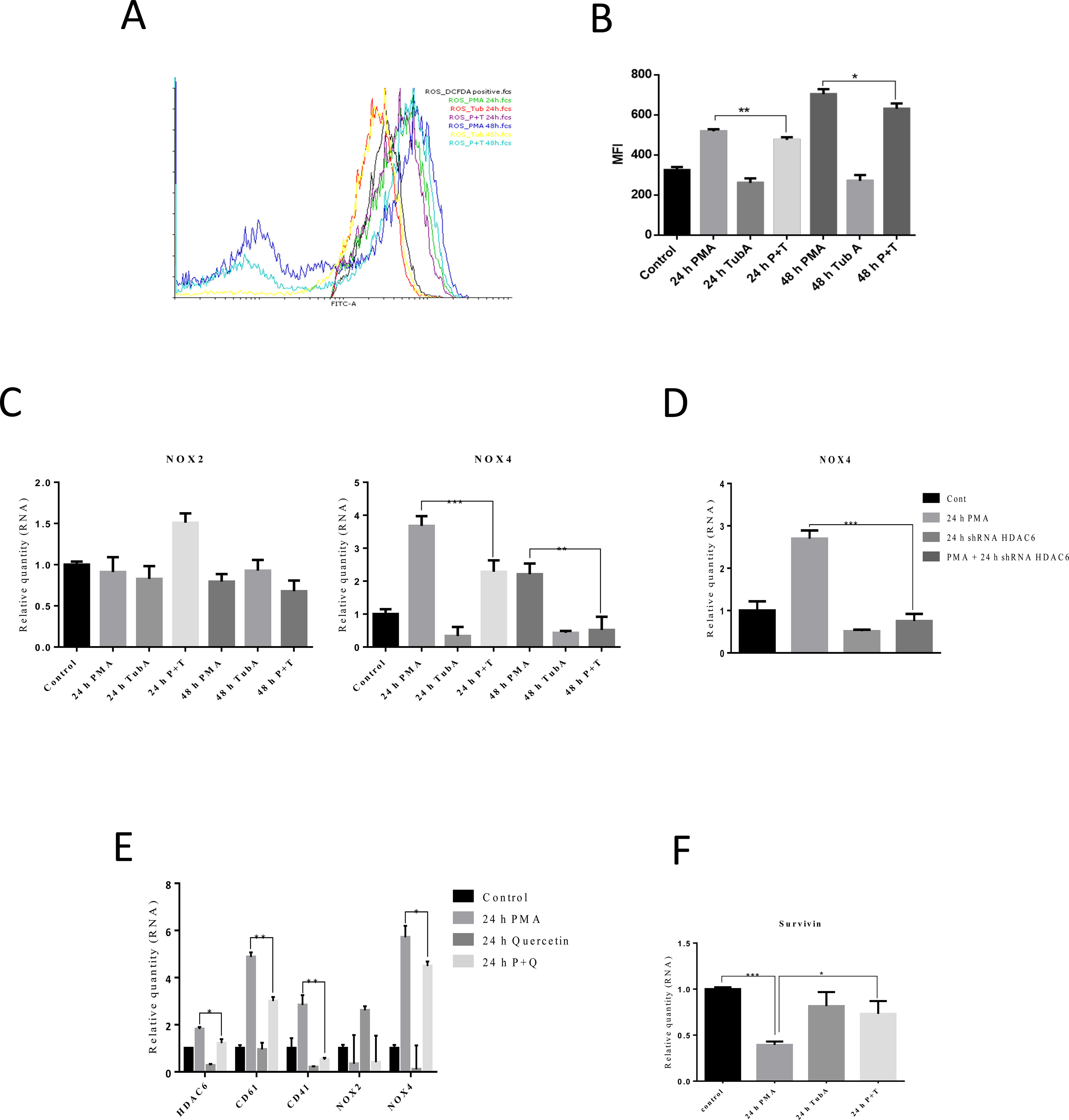
Crosstalk between PMA signalling, ROS and HDAC6 during MK differentiation. ROS measurement by FACS in PMA and / or Tubastatin A treated cells for 24 h and 48 h expressed as MFI (A) and overlay of histogram (A(i)), RNA expression analysis of NOX2, NOX4 during MK differentiation under HDAC6 inhibition (B), NOX4 expression in HDAC6 knockdown cells during MK differentiation (C), RNA expression analysis of HDAC6, MK markers, NOX2 and NOX4 in the presence of quercetin and or PMA (D), and RNA expression analysis of survivin in PMA and / or Tubastatin A treated cells for 24 h (E).

## Discussion

HDACs are involved in the regulation of gene expression and also control various biological processes by deacetylation of non-histone proteins ^27^. Hematopoiesis is a process of differentiation of blood cells from the hematopoietic stem cells that are regulated by various transcription factors, cytokines and miRNAs ^2^. Earlier studies have shown that sustained expression of HDAC1 by GATA1 transcription factor allows the differentiation of myeloid progenitor’s cells to erythrocyte and megakaryocytic lineages ^28^. Treatment of human primary CD34+ cells with sub-optimal concentration of panobinostat, a pan-HDAC inhibitor, does not affect the MK proliferation or commitment but inhibited maturation and platelet formation^29^. But the molecular mechanisms and specific roles of HDACs during the MK differentiation remain unclear. In this study we have shown the upregulation of Class IIb HDACs (HDAC6 and HDAC10) with decreased acetylation levels of H3K9 and H3K56 during PMA induced differentiation of K562 cells to megakaryocytes. HDAC6 is upregulated time dependently peaking at 4^th^ day of PMA treatment, and is required for the expression of MK markers CD61 and CD41, inhibition of erythroid lineage gene, GYPA and is involved in sustainable production of ROS.

HDAC6 is a class IIb cytoplasmic deacetylase with two homologous catalytic domains making it unique among all other HDACs ^30^. HDAC6 is involved in regulating the function stability, protein-protein or protein-DNA interaction, localization and signaling of cytoplasmic proteins by deacetylation, an important posttranslational modification. HDAC6 also possesses a zinc finger motif (ZnF-UBP) that enables it to bind with ubiquitin and thus plays a role in clearing the misfolded protein aggregates^31^. The well-studied cytoplasmic substrates of HDAC6 include tubulin, HSP90, cortactin and peroxiredoxins ^32-35^. In the nucleus, HDAC6 regulates expression of genes by interacting with nuclear proteins such as HDAC11, sumoylated p300, transcriptional repressors such as LCoR, ETO-2 ^36^ and transcription factors RUNX2 and NF-kappaB ^37 38^. Studies have shown the role of HDAC6 in variety of biological processes including metastasis, cell migration, autophagy, and viral infection^39,40 41^.

Our results in PMA induced K562 cell differentiation to MKs are in line with the recent studies by Messaoudi et al using CD34+ cells where the results have shown that HDAC6 is indispensable for proplatelet release from MKs by regulating the acetylation of cytoskeletal protein, cortactin ^25^. The study also demonstrated a significant decrease in MK cell number and Colony forming units (CFU) when CD34+ cells were treated with HDAC6 inhibitor (5µM Tubastatin A)^25^. Similarly our results demonstrate reduced MK marker expression in HDAC6 inhibited PMA-treated K562 cells. Therefore our results in the K562 cell line are biologically meaningful and hold significance as in CD34+ cells. Furthermore, Sardina JL et al. have demonstrated that the results obtained using K562 cells as megakaryocyte progenitor cells and the results obtained using CD34+ cells were same and reproducible suggesting the robustness of the PMA-induced MK differentiation of K562 cell line model system^42^.

Recent studies have shown that the nuclear HDAC6 promotes epithelial-mesenchymal transition (EMT) by suppressing the genes involved in formation of tight junctions ^43^. In our study also, we have observed the localization of HDAC6 in the nucleus predominantly, indicating that HDAC6 might be involved in suppression of erythroid lineage genes or regulating the activity or function of transcription factors. To further probe the role of nuclear HDAC6, we did ChIP-PCR for GYPA promoter. ChIP-PCR studies confirmed the binding sites of HDAC6 at GYPA promoter and the enrichment was increased in PMA-treated cells, significantly reduced in the presence of HDAC6 inhibitor. Interestingly we have observed the enrichment of HDAC6 to GYPA promoter in control cells (progenitor cells) also suggesting biasedness of MEP cells towards MK lineage due to the short half life of MK cells (5-7 days) compared to longevity of erythrocytes (120 days). However, further studies are needed to get a deeper insight into the ME-MK biasedness. HDACs are recruited to the target promoter regions by transcription factors and repressor proteins, as they do not contain DNA binding domain ^44^. Further studies are required to find out the interacting partner of HDAC6 in repressing GYPA. Thus in progenitor cells, GYPA is expressed at basal level due to the repressors bound to its promoter and repressed completely in the presence of lineage-specific cytokines or overexpression of lineage specific transcription factors.

At the molecular level, we have shown upregulation of HDAC6 *via* ERK1/2 signaling pathway and the down regulation of MK marker, CD61 when HDAC6 is inhibited or knockdown during MK differentiation. We have also observed increased expression of CD61 when HDAC6 is overexpressed in K562 cells giving us a clue that CD61 might be indirectly regulated by HDAC6. It is established that HDAC6 plays a role in ubiquitination and studies by ChonghuaLi, *et al*, have shown that SnoN, an inhibitor of Smad Pathway, is a repressor of CD61 and undergoes ubiquitination during PMA signaling of MK differentiation^45^. Furthermore our results are in line with ChonghuaLi, *et al* study showing upregulation of MK transcription factors GATA2 and FOG1. These results indicate presumably that HDAC6 might be the repressor of repressor (SnoN) of MK-specific transcription factors. Further studies are needed to confirm these presumptions.

Most of the studies have shown that ROS is required for MK cell commitment, maturation and for the release of platelets ^42,46^. ROS is the upstream regulator in activating the cascade of signaling proteins and finally leads to the activation of c-JUN, c-FOS, NF-κB transcription factors, which further regulate the expression of lineage specific genes. Studies have shown that treatment of cells with antioxidants (Trolox, NAC) inhibited the acquisition of MK morphological features and reduction in MK marker expression ^42^. p22phox-dependent NADPH oxidase is the primary source of ROS production ^42^ and the inhibition of ROS by antioxidants hinders MK differentiation from progenitor cells. But it is not clear, out of the four NOX homologues (NOX1-4), which NOX is responsible for the production of ROS. First time, our studies have shown that NOX4 is involved in PMA-induced ROS production. Youn GS *et al*, have shown that HDAC6 overexpression induces the ROS production by upregulating NADPH Oxidase in macrophages ^47^. In our study, we have observed downregulation of NOX4 when HDAC6 is inhibited during MK differentiation suggesting that HDAC6 plays important role in ROS production by regulating NOX4 expression. In astrocytes, crosstalk between HDAC6 and NOX2 regulates the expression of pro-inflammatory mediators during HIV-1 infection ^48^. In line with these results we found downregulation of HDAC6 and NOX4 when ROS is inhibited with an antioxidant, quercetin during MK differentiation indicating the positive feedback loop between HDAC6, NOX4 and ROS production. Zhang Y et al found that Aurora-B/AIM-1 and survivin, the 2 critical mitotic regulators were mislocalized or absent during endomitosis of mouse MKs^49^. This was further supported by finding by McCrann et al, who have shown that NOX4 is required for polyploidazation of vascular smooth muscle cells (VSMC) by down regulating survivin^50^. Our results have also shown, upregulation of survivin in the presence of HDAC6 inhibitor, where NOX4 expression was reduced.

In conclusion, our findings provide a new insight into the molecular mechanisms by which HDAC6 induces sustainable levels of ROS during MK differentiation by upregulating the NOX4 expression, positively regulating the expression of MK lineage transcription factors, GATA2 and FOG1, MK lineage marker CD61 and repressing erythroid lineage gene GYPA (Fig. 6).

**Fig 6:**
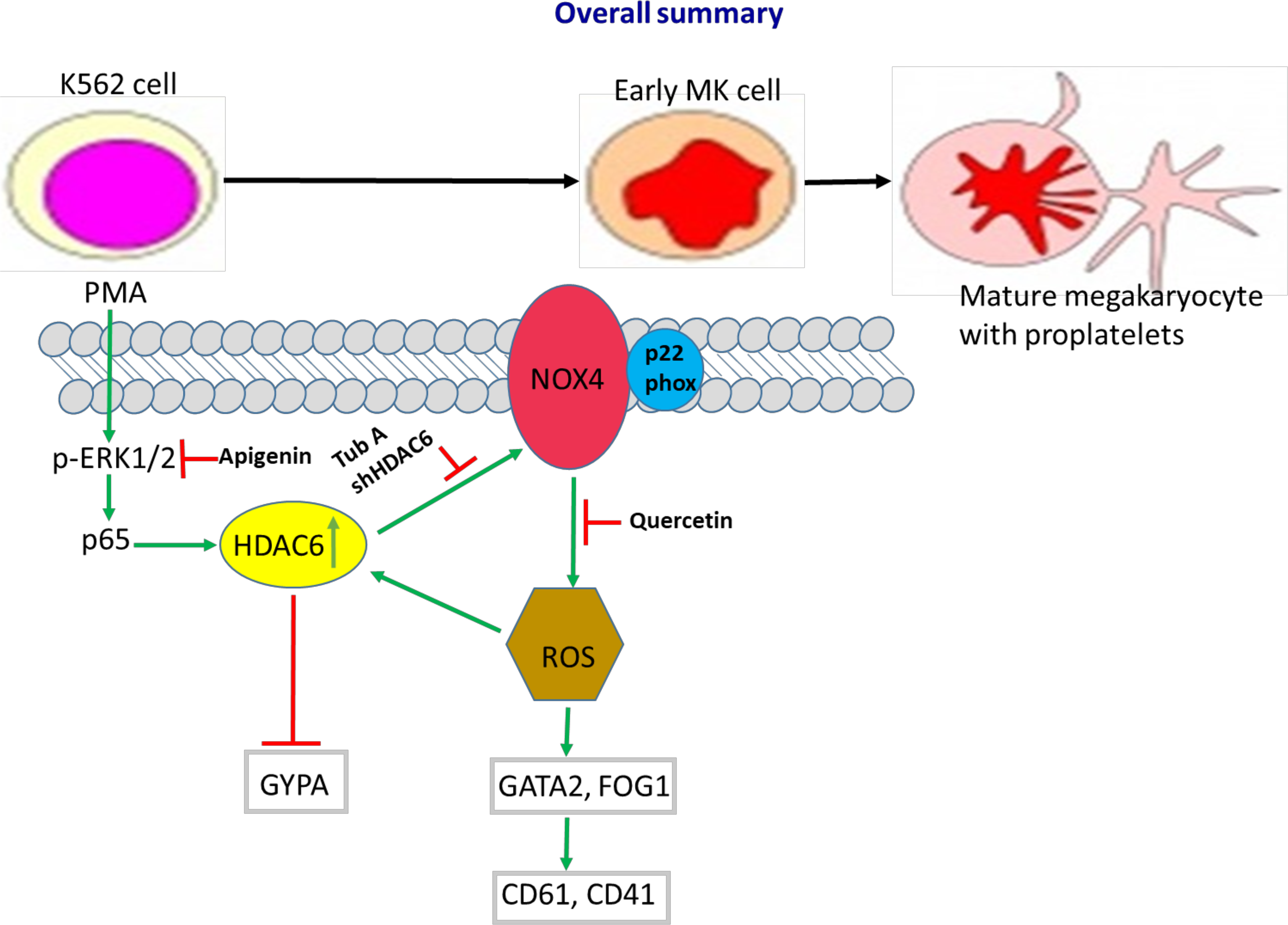
Schematic representation of observations from the study.

## Acknowledgements

Funding (in part) from CSIR (Grant# 37(1497)/11/EMR-II) and SERB (Grant# EMR/2015/001948) to AMK is highly acknowledged. Fellowship to GK from DBT (DBT-JRF/15/OM/621/3485) is acknowledged.

## Authorship Contributions

GK has performed the experiments, analyzed the results and drafted the manuscript. GRK gave critical suggestions throughout the study and reviewed the manuscript. AMK conceptualized the study, analyzed the results and reviewed the manuscript.

## Supplementary Information

**Fig. S1:** Immunoblot showing the HDAC6 protein levels in cytoplasmic (CE) and Nuclear extracts (NE) of control and PMA-treated K562 cells. HDAC6 levels in Total protein (TP) lysates is also included.

**Fig. S2:** Immunoblot showing the acetyl-tubulin levels of K562 cells treated with different concentrations of tubastatin A to determine the optimum concentration of Tubastatin A demonstrating HDAC6 inhibitory effect.

**Fig. S3:** Graphs showing the effect of tubastatin A on the expression (mRNA) of class I HDACs.

## References

1. Becker AJ, Mc CE, Till JE. Cytological demonstration of the clonal nature of spleen colonies derived from transplanted mouse marrow cells. Nature. 1963;197:452–454.

2. Orkin SH, Zon LI. Hematopoiesis: an evolving paradigm for stem cell biology. Cell. 2008;132(4):631–644.

3. Siminovitch L, McCulloch EA, Till JE. The Distribution of Colony-Forming Cells among Spleen Colonies. J Cell Comp Physiol. 1963;62:327–336.

4. Thomas D, Vadas M, Lopez A. Regulation of haematopoiesis by growth factors - emerging insights and therapies. Expert Opin Biol Ther. 2004;4(6):869–879.

5. Chen CZ, Li L, Lodish HF, Bartel DP. MicroRNAs modulate hematopoietic lineage differentiation. Science. 2004;303(5654):83–86.

6. Klimchenko O, Mori M, Distefano A, et al. A common bipotent progenitor generates the erythroid and megakaryocyte lineages in embryonic stem cell-derived primitive hematopoiesis. Blood. 2009;114(8):1506–1517.

7. Lindemann S, Tolley ND, Dixon DA, et al. Activated platelets mediate inflammatory signaling by regulated interleukin 1beta synthesis. J Cell Biol. 2001;154(3):485–490.

8. Suharti C, van Gorp EC, Setiati TE, et al. The role of cytokines in activation of coagulation and fibrinolysis in dengue shock syndrome. Thromb Haemost. 2002;87(1):42–46.

9. Ballem PJ, Segal GM, Stratton JR, Gernsheimer T, Adamson JW, Slichter SJ. Mechanisms of thrombocytopenia in chronic autoimmune thrombocytopenic purpura. Evidence of both impaired platelet production and increased platelet clearance. J Clin Invest. 1987;80(1):33–40.

10. Lozzio BB, Lozzio CB. Properties of the K562 cell line derived from a patient with chronic myeloid leukemia. Int J Cancer. 1977;19(1):136.

11. Shelly C, Petruzzelli L, Herrera R. PMA-induced phenotypic changes in K562 cells: MAPK-dependent and -independent events. Leukemia. 1998;12(12):1951–1961.

12. Villeval JL, Pelicci PG, Tabilio A, et al. Erythroid properties of K562 cells. Effect of hemin, butyrate and TPA induction. Exp Cell Res. 1983;146(2):428–435.

13. Butler TM, Ziemiecki A, Friis RR. Megakaryocytic differentiation of K562 cells is associated with changes in the cytoskeletal organization and the pattern of chromatographically distinct forms of phosphotyrosyl-specific protein phosphatases. Cancer Res. 1990;50(19):6323–6329.

14. Kang CD, Lee BK, Kim KW, Kim CM, Kim SH, Chung BS. Signaling mechanism of PMA-induced differentiation of K562 cells. Biochem Biophys Res Commun. 1996;221(1):95–100.

15. Murray NR, Baumgardner GP, Burns DJ, Fields AP. Protein kinase C isotypes in human erythroleukemia (K562) cell proliferation and differentiation. Evidence that beta II protein kinase C is required for proliferation. J Biol Chem. 1993;268(21):15847–15853.

16. Franklin CC, Kraft AS. Constitutively active MAP kinase kinase (MEK1) stimulates SAP kinase and c-Jun transcriptional activity in U937 human leukemic cells. Oncogene. 1995;11(11):2365–2374.

17. Kim KW, Kim SH, Lee EY, et al. Extracellular signal-regulated kinase/90-KDA ribosomal S6 kinase/nuclear factor-kappa B pathway mediates phorbol 12-myristate 13-acetate-induced megakaryocytic differentiation of K562 cells. J Biol Chem. 2001;276(16):13186–13191.

18. Chen S, Su Y, Wang J. ROS-mediated platelet generation: a microenvironment-dependent manner for megakaryocyte proliferation, differentiation, and maturation. Cell Death Dis. 2013;4:e722.

19. Cress WD, Seto E. Histone deacetylases, transcriptional control, and cancer. J Cell Physiol. 2000;184(1):1–16.

20. Eom GH, Kook H. Posttranslational modifications of histone deacetylases: implications for cardiovascular diseases. Pharmacol Ther. 2014;143(2):168–180.

21. de Ruijter AJ, van Gennip AH, Caron HN, Kemp S, van Kuilenburg AB. Histone deacetylases (HDACs): characterization of the classical HDAC family. Biochem J. 2003;370(Pt 3):737–749.

22. Gregoretti IV, Lee YM, Goodson HV. Molecular evolution of the histone deacetylase family: functional implications of phylogenetic analysis. J Mol Biol. 2004;338(1):17–31.

23. Walkinshaw DR, Yang XJ. Histone deacetylase inhibitors as novel anticancer therapeutics. Curr Oncol. 2008;15(5):237–243.

24. Kretsovali A, Hadjimichael C, Charmpilas N. Histone deacetylase inhibitors in cell pluripotency, differentiation, and reprogramming. Stem Cells Int. 2012;2012:184154.

25. Messaoudi K, Ali A, Ishaq R, et al. Critical role of the HDAC6-cortactin axis in human megakaryocyte maturation leading to a proplatelet-formation defect. Nat Commun. 2017;8(1):1786.

26. Shechter D, Dormann HL, Allis CD, Hake SB. Extraction, purification and analysis of histones. Nat Protoc. 2007;2(6):1445–1457.

27. Rundlett SE, Carmen AA, Kobayashi R, Bavykin S, Turner BM, Grunstein M. HDA1 and RPD3 are members of distinct yeast histone deacetylase complexes that regulate silencing and transcription. Proc Natl Acad Sci U S A. 1996;93(25):14503–14508.

28. Wada T, Kikuchi J, Nishimura N, Shimizu R, Kitamura T, Furukawa Y. Expression levels of histone deacetylases determine the cell fate of hematopoietic progenitors. J Biol Chem. 2009;284(44):30673–30683.

29. Iancu-Rubin C, Gajzer D, Mosoyan G, Feller F, Mascarenhas J, Hoffman R. Panobinostat (LBH589)-induced acetylation of tubulin impairs megakaryocyte maturation and platelet formation. Exp Hematol. 2012;40(7):564–574.

30. Zou H, Wu Y, Navre M, Sang BC. Characterization of the two catalytic domains in histone deacetylase 6. Biochem Biophys Res Commun. 2006;341(1):45–50.

31. Boyault C, Zhang Y, Fritah S, et al. HDAC6 controls major cell response pathways to cytotoxic accumulation of protein aggregates. Genes Dev. 2007;21(17):2172–2181.

32. Hubbert C, Guardiola A, Shao R, et al. HDAC6 is a microtubule-associated deacetylase. Nature. 2002;417(6887):455–458.

33. Rao R, Fiskus W, Yang Y, et al. HDAC6 inhibition enhances 17-AAG--mediated abrogation of hsp90 chaperone function in human leukemia cells. Blood. 2008;112(5):1886–1893.

34. Zhang X, Yuan Z, Zhang Y, et al. HDAC6 modulates cell motility by altering the acetylation level of cortactin. Mol Cell. 2007;27(2):197–213.

35. Parmigiani RB, Xu WS, Venta-Perez G, et al. HDAC6 is a specific deacetylase of peroxiredoxins and is involved in redox regulation. Proc Natl Acad Sci U S A. 2008;105(28):9633–9638.

36. Fernandes I, Bastien Y, Wai T, et al. Ligand-dependent nuclear receptor corepressor LCoR functions by histone deacetylase-dependent and -independent mechanisms. Mol Cell. 2003;11(1):139–150.

37. Zhang W, Kone BC. NF-kappaB inhibits transcription of the H(+)-K(+)-ATPase alpha(2)-subunit gene: role of histone deacetylases. Am J Physiol Renal Physiol. 2002;283(5):F904–911.

38. Westendorf JJ, Zaidi SK, Cascino JE, et al. Runx2 (Cbfa1, AML-3) interacts with histone deacetylase 6 and represses the p21(CIP1/WAF1) promoter. Mol Cell Biol. 2002;22(22):7982–7992.

39. Lee YS, Lim KH, Guo X, et al. The cytoplasmic deacetylase HDAC6 is required for efficient oncogenic tumorigenesis. Cancer Res. 2008;68(18):7561–7569.

40. Lee JY, Koga H, Kawaguchi Y, et al. HDAC6 controls autophagosome maturation essential for ubiquitin-selective quality-control autophagy. EMBO J. 2010;29(5):969–980.

41. Zhang L, Ogden A, Aneja R, Zhou J. Diverse roles of HDAC6 in viral infection: Implications for antiviral therapy. Pharmacol Ther. 2016;164:120–125.

42. Sardina JL, Lopez-Ruano G, Sanchez-Abarca LI, et al. p22phox-dependent NADPH oxidase activity is required for megakaryocytic differentiation. Cell Death Differ. 2010;17(12):1842–1854.

43. Ding G, Liu HD, Huang Q, et al. HDAC6 promotes hepatocellular carcinoma progression by inhibiting P53 transcriptional activity. FEBS Lett. 2013;587(7):880–886.

44. Shahbazian MD, Grunstein M. Functions of site-specific histone acetylation and deacetylation. Annu Rev Biochem. 2007;76:75–100.

45. Li C, Peart N, Xuan Z, Lewis DE, Xia Y, Jin J. PMA induces SnoN proteolysis and CD61 expression through an autocrine mechanism. Cell Signal. 2014;26(7):1369–1378.

46. Mostafa SS, Miller WM, Papoutsakis ET. Oxygen tension influences the differentiation, maturation and apoptosis of human megakaryocytes. Br J Haematol. 2000;111(3):879–889.

47. Youn GS, Lee KW, Choi SY, Park J. Overexpression of HDAC6 induces pro-inflammatory responses by regulating ROS-MAPK-NF-kappaB/AP-1 signaling pathways in macrophages. Free Radic Biol Med. 2016;97:14–23.

48. Youn GS, Cho H, Kim D, Choi SY, Park J. Crosstalk between HDAC6 and Nox2-based NADPH oxidase mediates HIV-1 Tat-induced pro-inflammatory responses in astrocytes. Redox Biol. 2017;12:978–986.

49. Zhang Y, Nagata Y, Yu G, et al. Aberrant quantity and localization of Aurora-B/AIM-1 and survivin during megakaryocyte polyploidization and the consequences of Aurora-B/AIM-1-deregulated expression. Blood. 2004;103(10):3717–3726.

50. McCrann DJ, Yang D, Chen H, Carroll S, Ravid K. Upregulation of Nox4 in the aging vasculature and its association with smooth muscle cell polyploidy. Cell Cycle. 2009;8(6):902–908.

