## Supplementary figures and images for "HDAC6 promotes PMA-induced megakaryocyte differentiation of K562 cells by regulating ROS levels *via* NOX4 and repressing Glycophorin A"

### Supplementary Fig

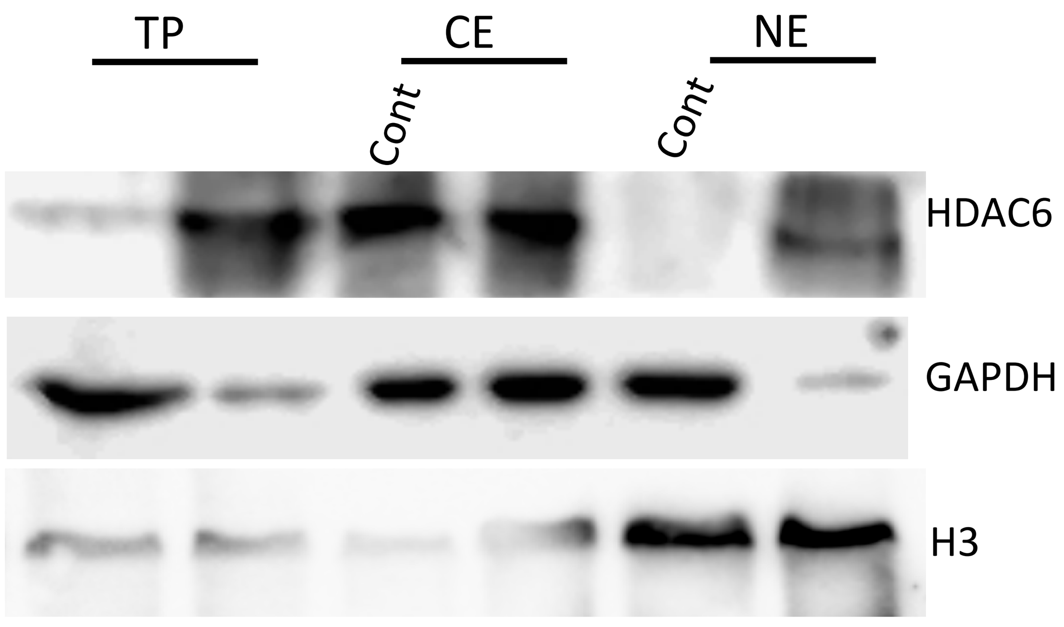

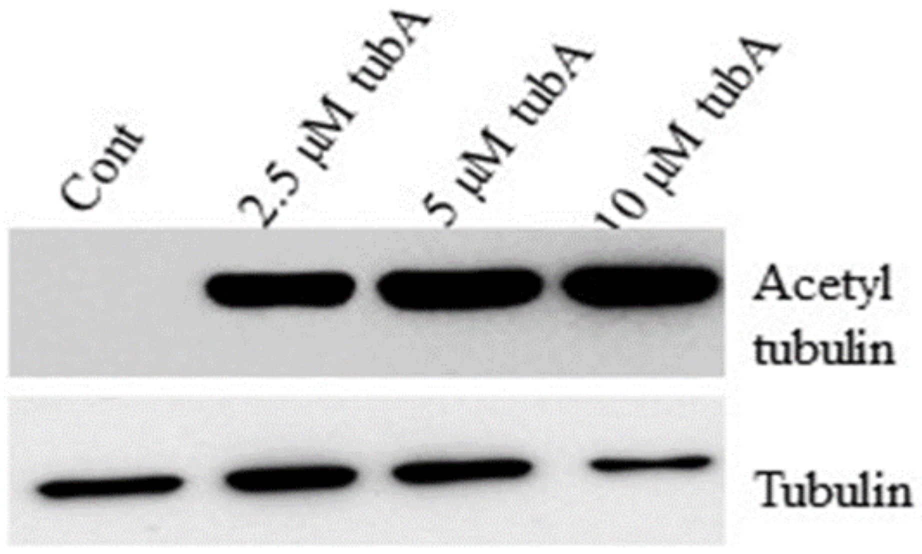

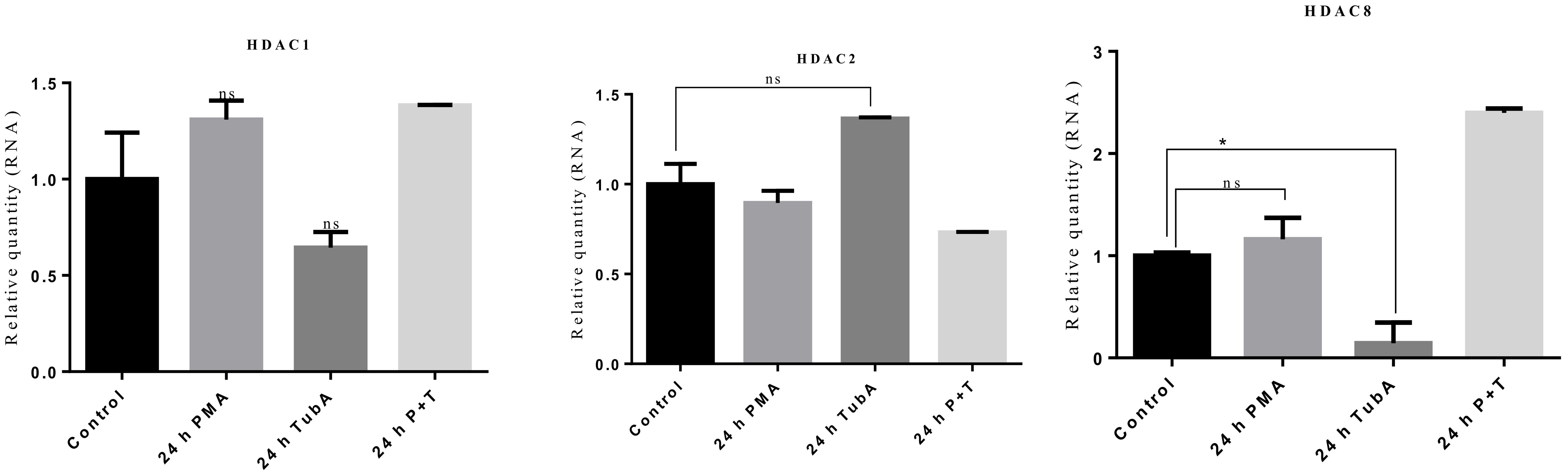
